# The detection of brood parasitism and quasi-parasitism in the burying beetle *Nicrophorus quadripunctatus* under natural conditions

**DOI:** 10.1101/2023.01.11.523691

**Authors:** Takuma Niida, Izumi Yao, Tomoyosi Nisimura, Seizi Suzuki

## Abstract

Intraspecific brood parasitism (IBP), where a parasitic female lays eggs in the nest of another female of the same species, occurs in insects and birds. Also, quasi-parasitism (QP), where a parasitic female copulates with a host male at his nest and lays eggs that are fertilized by the male, has been documented in a few monogamous birds, but QP has not been observed in any insects.

Burying beetles, genus *Nicrophorus*, use small vertebrate carcasses for reproducing and providing biparental care for their offspring. IBP has been observed in one burying beetle by laboratory experiments, but has not been well reported under natural conditions. IBP and QP may occur under natural conditions in burying beetles.

Here we focused on a burying beetle, *Nicrophorus quadripunctatus*. Ten broods, consisting of larvae and their parental female and male, were collected from a deciduous forest. To investigate the kin relationship between parents and larvae, eight microsatellite DNA loci were used.

We detected three types of parasitic larvae: 1) larva not related to either its parental female or male, 2) larva not related to its parental female, but unknown regarding its parental male, and 3) larva not related to its parental female, but related to its parental male. These results suggested that IBP and QP can occur with certain frequencies in the reproduction of *N. quadripunctatus* under natural conditions. QP is thought to have a benefit for a parental male to enhance his paternity within one brood in this species.

## Introduction

In many species which provide parental care for a brood in reproduction, some females lay eggs into a nest of another female of the same species. This type of parasitism for parental care had been roughly called intraspecific brood parasitism (IBP) before in studies in which microsatellite DNA analysis was commonly used (Griffith et al., 2004). However, by performing DNA analysis of kin relationships, it has been found that various types of the parasitism exist depending on the relationships among a host, parasite, and brood. For example, using such analysis, we are able to discriminate the relationship between offspring and a parental female which is not a genetic parent as extra-pair maternity (EPM) (Akçay & Roughgarden, 2007). For a parental male, the relationship is called extra-pair paternity (EPP). Thus, currently, the parasitism is designated as IBP when both EPM and EPP are detected in the nest where a female and male couple provides parental care, and when EPM is detected in the nest where a single female provides parental care (Tallamy, 2005; Akçay & Roughgarden, 2007).

In addition, microsatellite DNA analysis is able to discriminate the parasitism in which only EPM is detected but EPP is not detected in the nest where a female and male couple provides parental care (Akçay & Roughgarden, 2007). This type of parasitism is called quasi-parasitism (QP), which would occur when a female copulates with a host male at his nest and lays eggs that are fertilized by the male. QP has been reported in a few birds (Griffith et al., 2004; Li et al., 2009; Petrželková et al., 2015), but not in insects, though IBP has been found commonly in birds and insects (Heteroptera, Coleoptera and Hymenoptera) (Yom-Tov, 2001; Tallamy, 2005; Lyon & Eadie, 2008).

Burying beetles, genus *Nicrophorus* (Coleoptera: Silphidae: Nicrophorinae), use small vertebrate carcasses to prepare for parental care (Scott, 1998). To monopolize the carcass, aggressive intrasexual competition occurs, where larger adults win in almost all cases (Bartlett & Ashworth, 1988; Trumbo, 1990). The winning female lays eggs in the soil surrounding the carcass, and then the female or the female and male couple provides parental care for hatching larvae by regurgitating (Scott, 1998).

Meanwhile, the losing female has a chance to lay her eggs near the carcass because there is no physical barrier around the carcass. Indeed, parasitic behavior, namely IBP, has been demonstrated in *Nicrophorus vespilloides* under laboratory experiments (Müller et al., 1990; Eggert & Müller, 2011; Richardson et al., 2021).

However, few studies about IBP exist under natural conditions and there is no report other than one about *N. vespilloides* (Müller et al., 2007). Additionally, in this species, a parental male is considered to sometimes mate with another female, e.g. a parasitic female (Alves & Bryant, 1998; Houston et al., 2005; Paquet & Smiseth, 2016), so that QP also could occur in the reproduction of *Nicrophorus*.

Here we focused on the reproductive behavior in a natural population of *Nicrophorus quadripunctatus* (Kraat, 1877), which is widely distributed in Japan, and examined kin relationship between parents and offspring in their broods, using microsatellite DNA markers (Sikes & Venables, 2013; Suzuki & Yao, 2014). The purposes of the present study were to detect the presence of larvae unrelated to parental females under natural conditions, and if present, to examine whether the parasitic larvae are derived from IBP or QP.

## Materials and methods

### Sample collection

*Nicrophorus quadripunctatus* broods were collected from a deciduous forest near the campus of Nihon University College of Bioresource Sciences (NUBS) in Fujisawa City, Japan (35°22’N, 139°27’E) (Figure 1; Supplementary data 1). A piece of chicken bait (approximately 25 g) was put on soil in an earthenware flowerpot as a food resource for reproduction of *N. quadripunctatus*. We set 66 pots in the field. When *N. quadripunctatus* adults were observed to have buried the chicken bait and raised larvae, the adults and larvae were referred to as one brood. A brood was preserved in a 50-mL conical tube with 99.9% ethanol.

**Figure 1.**
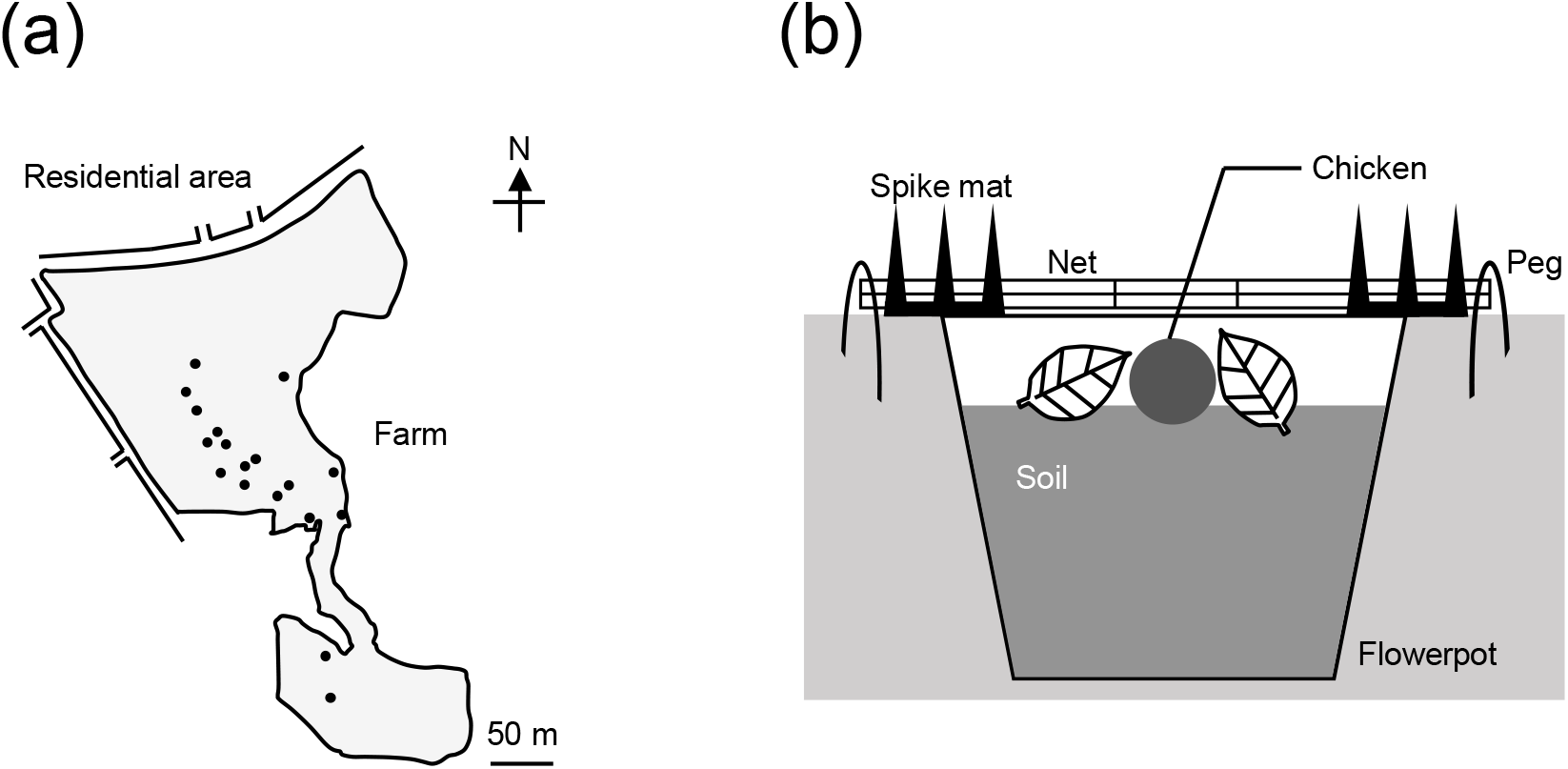
Reproductive situation of *Nicrophorus quadripunctatus* under natural conditions in this study. (a) General map of study area. Grey area represents a deciduous forest near NUBS. Black points represent 18 collection sites. (b) Lateral view of an earthenware flowerpot.

### Molecular experiments

DNA was extracted from larvae, adult females and adult males. Eight microsatellite DNA loci (Nq-01, -03, -04, -05, -07, -08, -09 and -10) were amplified (Suzuki & Yao, 2014). Then, to determine the fragment size of the PCR product, polyacrylamide gel electrophoresis was carried out with loading of the mixture of PCR product and DNA size standard marker into each well.

### Parentage analyses

The numbers and fragment size of alleles were determined using the gel image analyzing function of ImageJ (Schneider et al., 2012). When the fragment size of the target PCR product in larvae was ≥ 2 bp different from that in their parental female, it was regarded that the larvae had a different allele, based on the repeat motifs of the microsatellite DNA sequences (Suzuki & Yao, 2014).

The mismatch distribution between parental females and the larvae in each brood was checked for all eight loci. When a parental female and her larvae did not share at least one allele at any given locus, it was regarded that the larvae had a mismatched locus.

Subsequently, allele frequency analyses were performed for the alleles of adults using CERVUS (Marshall et al., 1998; Kalinowski et al., 2007; Koch et al., 2008; Jones et al., 2010; Flanagan & Jones, 2019). Parentage analyses were also performed using CERVUS for all eight loci. First, maternity analysis was conducted for the alleles of larvae and parental females (for parameter settings refer to Supporting information). Second, paternity analysis was conducted for the alleles of larvae unassigned to their parental females and parental males.

## Results

Thirteen pots were observed to carry *N. quadripunctatus* broods and collected. Three broods were excluded from molecular analyses due to absence of a parental female or too small brood size (Supplementary data 2). Thus, genotyping and parentage analyses were performed on four broods where one adult pair and larvae were found in each brood (broods: A, B, C and D), and six broods where one female and larvae were found in each brood (broods: E, F, G, H, I and J) (Figure 2).

**Figure 2.**
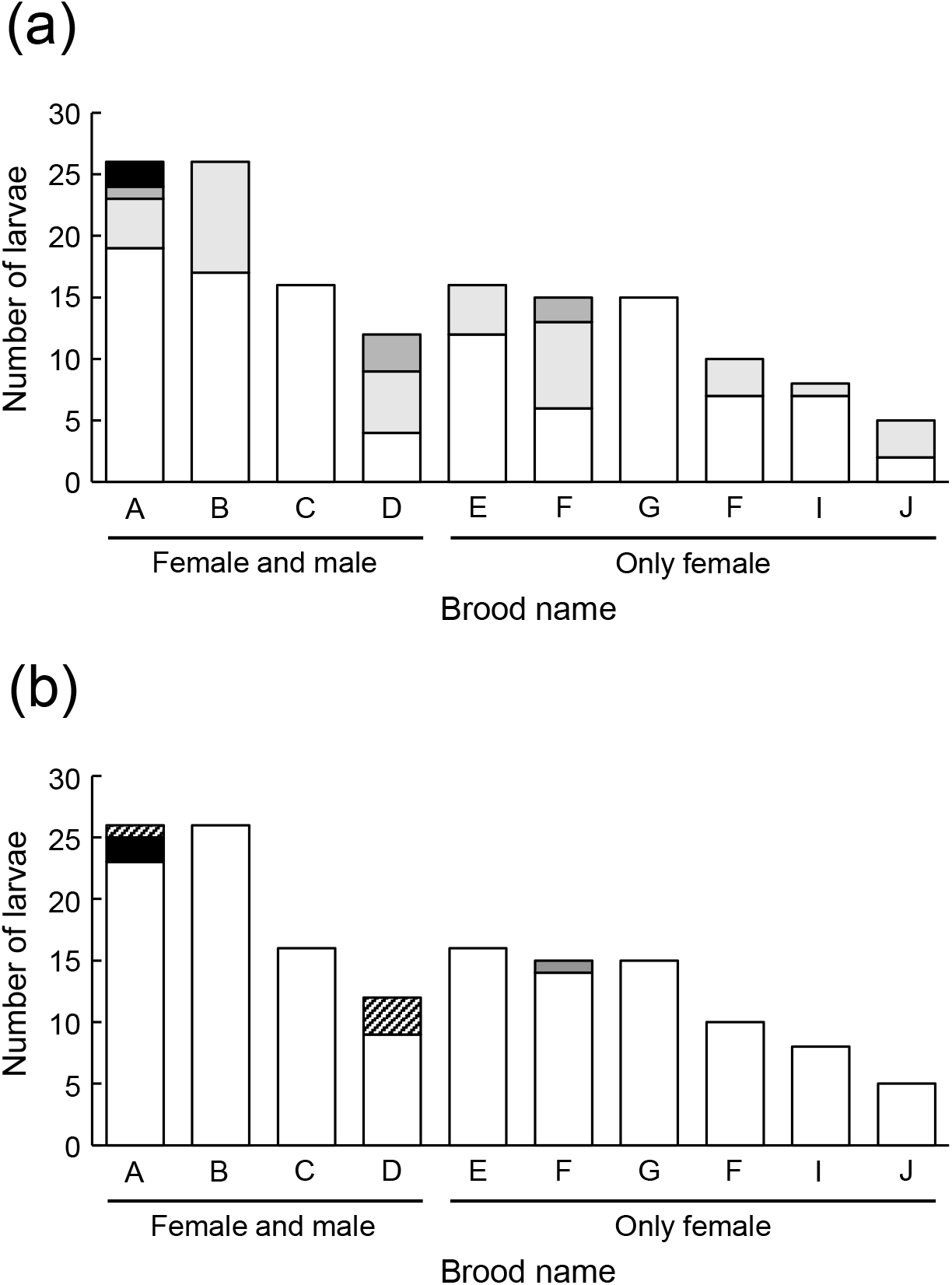
Analyses using eight microsatellite DNA loci in the broods of *Nicrophorus quadripunctatus*. (a) The mismatch distribution between parental females and their larvae in each brood. White, pale grey, grey and black bar represent the larvae mismatched at zero, one, two and three loci, respectively. (b) Parentage analysis between parents and their larvae in each brood, estimated by CERVUS. White bars represent the larvae assigned to parental females. In the broods with parental males (broods: A, B, C and D), black bar represents the larvae not assigned to both a parental female and a parental male, and hatched bars represent the larvae not assigned to parental females but assigned to parental males. In the broods without parental males (broods: E, F, G, H, I and J), grey bar represents the larva not assigned to a parental female.

Regarding the mismatch distribution between parental females and their larvae in each brood, 105 larvae were matched to their respective parental females. On the other hand, 36, 6 and 2 larvae were mismatched to their respective parental females at 1, 2 and 3 loci, respectively (Figure 2a). For the parentage analyses using CERVUS, 142 larvae were assigned to their respective parental females at 80% and 95% confidence level, and seven larvae were not assigned to their respective parental females (Figure 2b). Of these seven larvae, two larvae in one brood (brood A) were not assigned to the respective parental male, and for one larva in one brood (brood F) it was unclear whether it was assigned to the parental male due to the absence of a male. And four larvae in two broods (one larva in brood A, three larvae in brood D) were assigned to the respective parental males at 80% and 95% confidence level.

CERVUS detected two larvae unrelated to both their respective parental female and parental male, i.e., were designated EPM and EPP, in one brood (brood A; Figure 2b). Also, it detected one larva which was unrelated to its parental female, and which was thus designated EPM, in one brood (brood F; Figure 2b). When takeover of a resource happens by a subsequently coming female in one nest, offspring unrelated to a parental female may be derived from a previously reproducing female in the nest, in addition to being derived from a parasitic female (Waldeck & Andersson, 2006). In burying beetles, however, larvae and eggs produced by a previously reproducing female are cannibalized by a subsequently coming female (Scott, 1997; Eggert & Müller, 2011; Richardson et al., 2021). According to this finding from previous studies, the three larvae unrelated to parental females in this study were more likely derived from parasitic females, rather than previously reproducing females. Thus, intraspecific brood parasitism (IBP) can occur in *N. quadripunctatus* reproductive behavior.

In addition, CERVUS showed that four larvae were not related to their respective parental females but were related to their respective parental males, which means that only EPM and not EPP, occurred in two broods (brood A and D; Figure 2b). In the competition of burying beetles, losing females do not immediately depart from the carcass (Müller et al, 1990). They can copulate with the winning male and oviposit eggs fertilized by the male, resulting in the existence of larvae related to only the parental male. This result suggests that quasi-parasitism (QP) occurs in *N. quadripunctatus* reproductive behavior.

However, Griffith (2004) claimed that the occurrence of QP is far more limited that generally believed. For example, in burying beetles, it is known that a parental male tends to exclude an intruding female with the cooperation of a parental female (Otronen, 1988; Suzuki, 2011). Moreover, it was demonstrated that a parental male was manipulated by its parental female’s chemical substance, methyl geranate, which acts on the parental male as an anti-aphrodisiac to inhibit copulation with other females (Eggert & Sakaluk, 1995; Engel et al., 2016). These previous studies suggest that copulation of a parental male with a parasitic female is difficult, once one male and female become a reproductive pair.

Nevertheless, it is likely that there is a benefit for a parental male to mate with a parasitic female. A parental female generally copulates with multiple males, so that several larvae are derived from other than a parental male, decreasing his paternity (Pascoal et al., 2018). Against that, a parental male can copulate with a parasitic female, because the parasitic female is expected to oviposit eggs fertilized with the male into his nest, enhancing his paternity (Alves & Bryant, 1998). Because of this parental male’s advantage, QP can have evolved in *N. quadripunctatus*.

In summary, this study shows that IBP can occur with a certain frequency in broods of *N. quadripunctatus* under natural conditions. Furthermore, it suggests that QP exists in *N. quadripunctatus*, as well as in monogamous birds. In future studies, mechanisms for establishing QP, for example male’s pheromone emission to copulate with parasitic females (Müller & Egger, 1987), should be examined.

## Acknowledgements

We would like to thank Hiroshi Abé, Hirohiko Takeuchi, Hidetoshi Iwano, Yoshinori Hatakeyama and Yuuichi Yamamoto of College of Bioresource Sciences, Nihon University, and Shigeyuki Koshikawa of Faculty of Environmental Earth Science, Hokkaido University, for providing the opportunity to conduct this work; Kengo Noma, Nagisa Tosano, Kohei Seto, Masafumi Hasegawa, Mikio Kasatani, Natsumi Katsube for discussion and sample collection; and Elizabeth Nakajima for English editing. This study was supported by KAKENHI (Grant No. 20K06808).

## Author contributions

Seizi Suzuki, Tomoyosi Nisimura and Takuma Niida designed the experiments. Takuma Niida and Izumi Yao carried out the experiments and analyzed the data. All authors contributed to the writing.

## Data availability statement

The data that support the findings of this study are available on request from the corresponding author. The data are not publicly available due to privacy or ethical restrictions.

## Supporting information

Additional supporting information may be found online in the Supporting Information section at the end of the article.

---

**Supplementary data 1.**
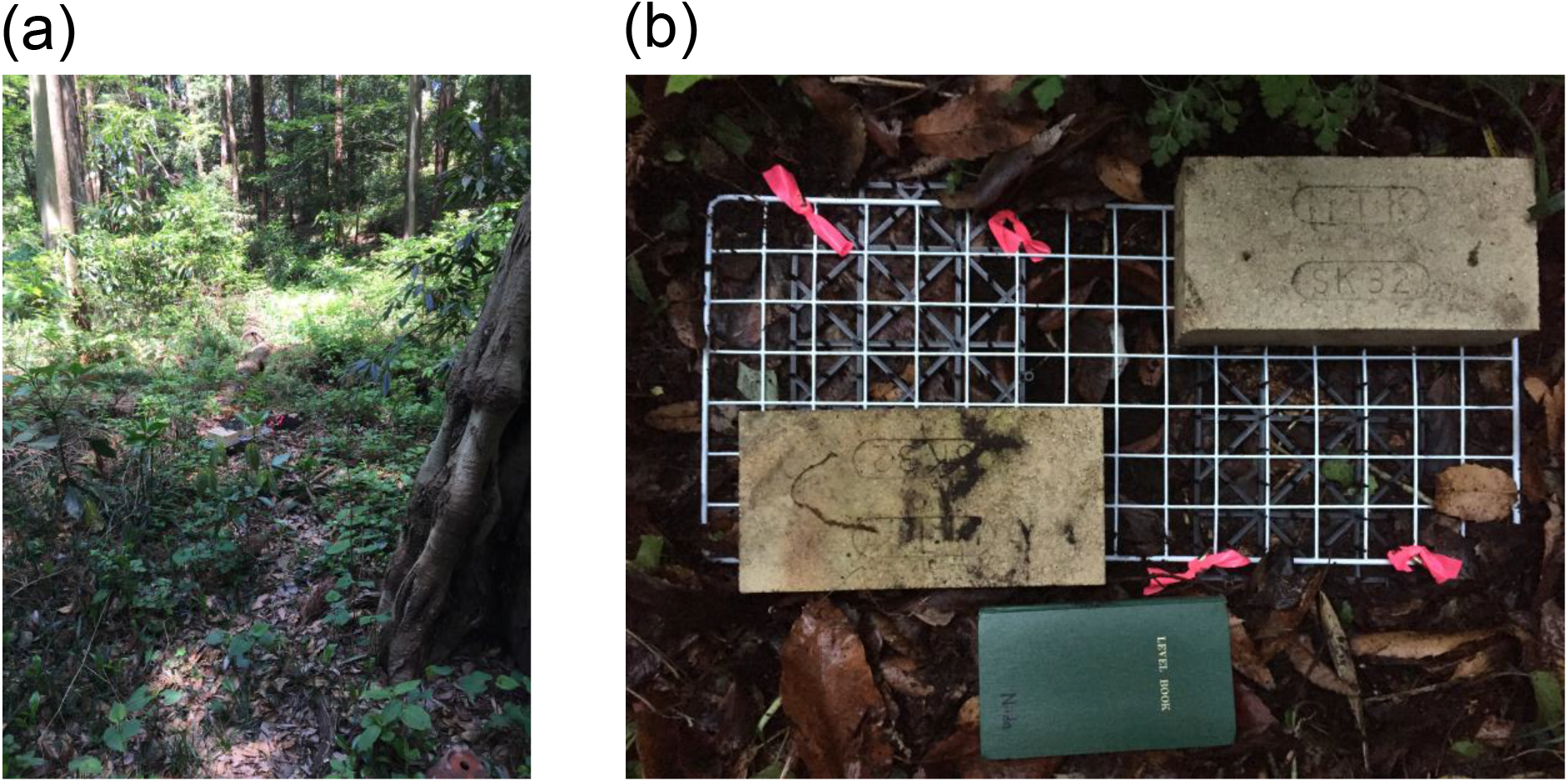

**Supplementary data 2.**
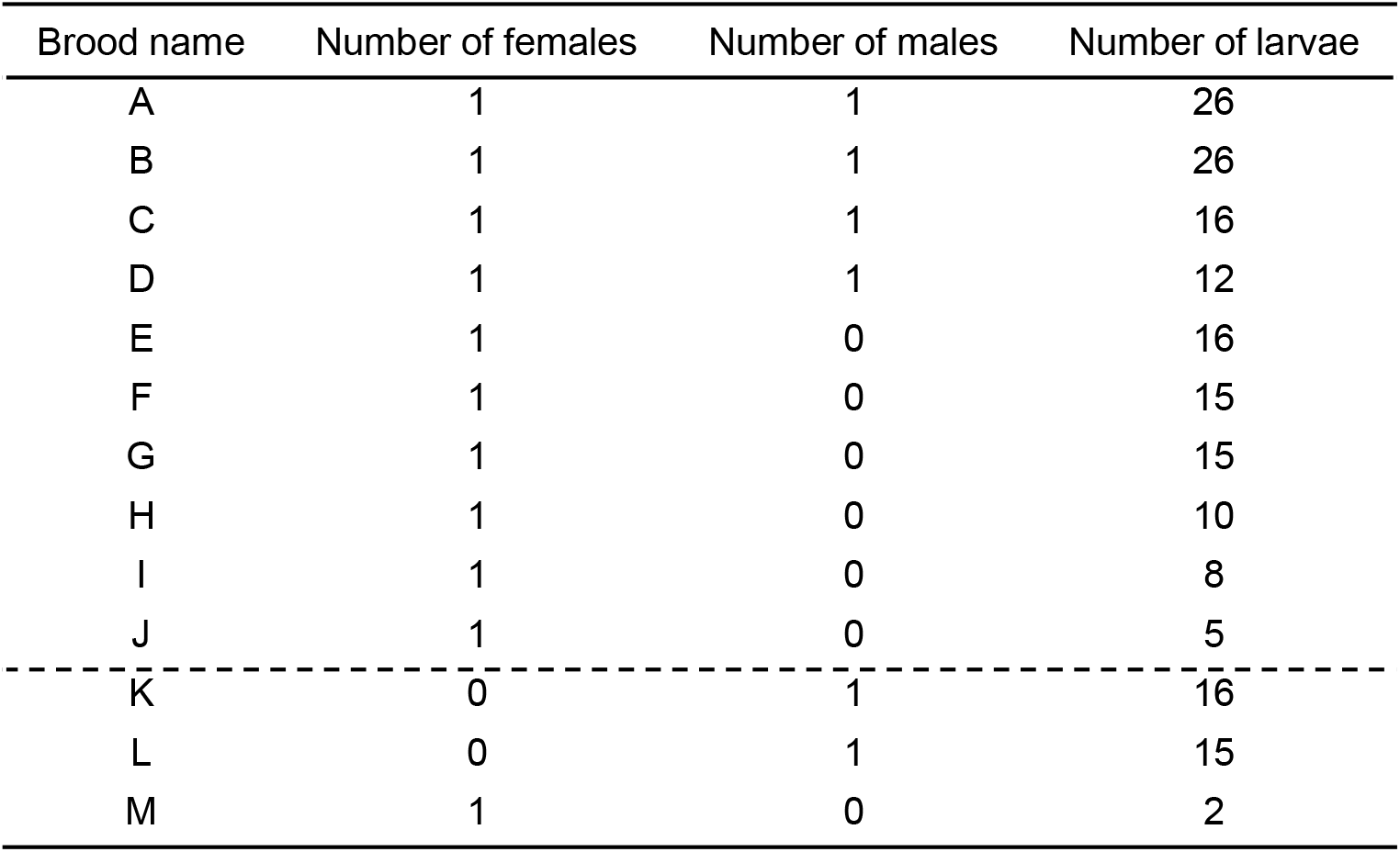

**Supplementary data 3.**
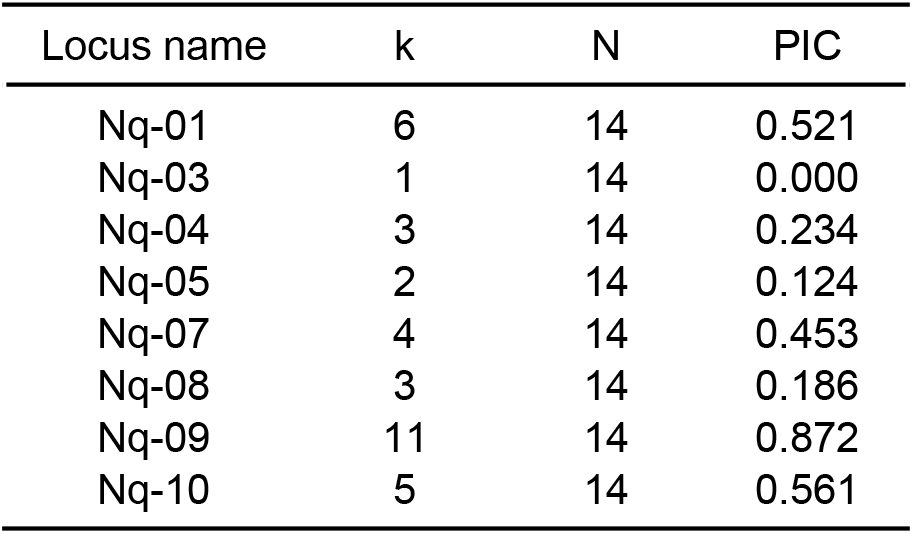

