## supplemental information for "The detection of brood parasitism and quasi-parasitism in the burying beetle *Nicrophorus quadripunctatus* under natural conditions"

Supporting information

Materials and methods

Study area and the installation of chicken

All broods of *Nicrophorus quadripunctatus* used for molecular work were collected from a deciduous forest near the campus of Nihon University College of Bioresource Sciences (NUBS) in Fujisawa city, Japan (35°22´N, 139°27´E) from April to May and from September to October in 2018 (Supplementary data 1a). Earthenware flowerpots (mostly flowerpots: 16 and 10 cm in diameter for top and bottom, 12 cm height) were placed into the ground so that their opening was at the level of the ground surface, and were set at more than 10-m intervals (Figure 1a). At each collection site, a flowerpot was filled with the forest soil up to about 3 cm from the top. A piece of raw chicken (ca. 25 g) was put on the soil in the flowerpot (Figure 1b; Supplementary data 1b) as a reproductive resource. To avoid disturbance by vertebrate scavengers, the flowerpot was covered by a sturdy steel net (mesh size 3.6 cm × 3.4 cm) and spike mats, which were fixed in the ground with pegs and bricks.

Collection of broods

We checked the pods in the afternoon about 3 three days to examine whether *N. quadripunctatus* adults started to reproduce. When *N. quadripunctatus* adults were observed to have buried the chicken and raised hatching larvae in a pot, the adults and larvae were referred as one brood and preserved in a 50 mL-conical tube with 99.9% ethanol. The tubes were stored at 5 ^O^C until molecular work. The broods with fewer than three larvae and those without a parental female were not used for the analyses (Supplementary data 2).

DNA extraction and PCR

Larvae that were larger than around 5 mm in body length were cut at the head (ca. 2-3 mm) with a disposable blade and it was used for DNA extraction. For those less than 5 mm in body length, the whole body was crushed. For adults, six legs were cut from the body and used for DNA extraction. DNA was extracted using the Wizard Genomic DNA Purification Kit (Promega, Tokyo) and Proteinase K solution (Kanto Chemical, Tokyo) according to the manufacturer’s instructions. Because the DNA concentration in some extracted DNA samples was too high to amplify the target regions, those DNA samples were diluted 100 fold before amplification. Eight microsatellite DNA loci (Nq-01, -03, -04, -05, -07, -08, -09, and -10) (Suzuki & Yao, 2014) were amplified using a thermal cycler T-100 (Bio-Rad, Tokyo). The PCR reaction solution was 10 µL in volume including 2 µL of 5 × KapaTaq Extra buffer (Nippon Genetics, Tokyo), 1 µL of 25 mM MgCl_2_ (final conc. 2.5 mM), 0.3 µL of 10 mM dNTP, 0.5 µL of forward and reverse 10 pM primers (final conc. 0.5 pM), 0.05 µL of KapaTaq Extra DNA polymerase, 4.65 µL of sterilized water, and 1 µL of DNA solution. The reaction cycle parameters were as follows: 1 cycle of first denaturation at 94 ^O^C (1 min.), 40 cycles of denaturation at 94 ^O^C (15 sec.), annealing of primers at 55 ^O^C (for every primer, 20 sec.), and extension at 68 ^O^C (1 min.), followed by 1cycle of final extension at 68 ^O^C (1 min.).

Polyacrylamide gel electrophoresis

To determine allele size of each locus, polyacrylamide gel electrophoresis (PAGE) was carried out. In this study, PAGE was run by loading the mixture of PCR product and DNA size standard marker into each well in order to determine the size of PCR product using DNA size standard marker in each lane. The PAGE protocol was as follows: 6 µL of sample solution included 1 µL of OneSTEP Marker 5 (φX174/HincII digest) (NIPPON GENE, Tokyo), 1 µL of PCR product amplified with Nq-04 or Nq-09 primers, 1 µL of Xylene-Cyanol loading dye buffer, and 3 µL of sterilized water. For PCR product amplified with other primers, OneSTEP Marker 9 (φX174/HinfI digest) was used as DNA size standard marker. PAGE was performed using an AE-6220 Dual Slab Chamber (ATTO, Tokyo). The sample solution was electrophoresed on a 13-cm nondenaturing gel consisting of 4.4 mL of 40% (final conc. 8%) acrylamide solution (29:1 acrylamide/bisacrylamide) (Nacalai tesque, Kyoto), 4.5 mL of 5 × TAE buffer, 155 µL of 10% ammonium persulphate solution (Wako, Tokyo), 20 µL of N,N,N,N-Tetramethylethylene-Diamine (TEMED) (MP Biomedicals, Tokyo), and 13.2 mL of distilled water. The electrophoresis was run at 200 V during 2.5 hours. Gels were stained with ethidium bromide and digital pictures were taken using an ultraviolet transilluminator, Printgraph AE-6932GX (ATTO, Tokyo).

Genotyping

The numbers and sizes of alleles at each locus were determined using the gel image analyzing function implemented in the software ImageJ 1.52q (Schneider et al., 2012). A series of DNA fragments, including the target PCR product bands and DNA size standard marker bands (335, 297, 291, 210, and 162 bps of Marker 5; 311, 249, 200, 151, and 140 bps of Marker 9), in a lane on a gel image was selected and converted to distinct sharp peaks on a lateral direction chart with X- and Y-axes. The values of X-axis were read from the points on the top of each of the peaks, indicating the migration of DNA fragments from an arbitrary selected origin (335 bp of Marker 5 and 311 bp of Marker 9 were set at zero as origin). A scatter plot, containing multiple points of the sharp peaks of DNA size standard marker and target PCR product, was created and subsequently the linear approximation along the multiple points was produced using Microsoft Excel. The allele sizes were determined by putting the X-axis value of a target PCR product into the linear approximation. Some non-specific DNA fragments that appeared at the same position in different lanes were also used to identify alleles. When the size of the target PCR product in larvae was ≥ 2 bp different from that in their parental female, it was regarded as a different allele, based on the repeat motifs of the microsatellite DNA sequences (Suzuki & Yao, 2014).

Parentage analyses

The mismatch distribution between parental females and their larvae in each brood was checked. When a parental female and her larvae did not share at least one allele at any given locus, it was regarded that the larvae had a mismatched locus.

Allele frequency analyses were performed with CERVUS version 3.0.7 (Marshall et al., 1998; Kalinowski et al., 2007). The number of alleles and polymorphic information content (PIC) at each locus were checked for parental females and males (Supplementary data 3) (Jones et al., 2010; Flanagan & Jones, 2019).

Parentage analyses were also performed with CERVUS. In this step, CERVUS calculates a log-likelihood (LOD) score as the natural logarithm of the ratio between the likelihood that the candidate parent is the true parent and the likelihood that the candidate parent is not the true parent (Koch et al., 2008; See www.fieldgenetics.com). Firstly, maternity analysis was conducted on the set of 10 parental females and 149 larvae in 10 broods (See Results and Discussion). In the simulation of the maternity analysis, parameters were set as 10,000 offspring (default) were produced by two candidate mothers with 50% parents sampled, 100% proportion of loci genotyped, 1% genotyping error rate (default), four minimum typed loci (recommended number) and confidence levels assessed by LOD distribution (default: relaxed > 80%, strict > 95%). Secondly, paternity analysis was conducted on the set of the larvae unassigned to any parental females and parental males (See Results and Discussion). Parameters were set to the same as those of the maternity analysis.

Legends

Supplementary data 1

(a) Landscape near collection site in a deciduous forest of NUBS. (b) Photograph of an earthenware flowerpot taken from above.

Supplementary data 2

Collected broods of *Nicrophorus quadripunctatus*. Three broods (K, L and M) below dashed line were excluded from molecular experiments.

Supplementary data 3

Summary statistics of eight microsatellite loci using 14 adults of *Nicrophorus quadripunctatus*. k, N and PIC represent number of alleles detected, the number of adults and polymorphic information content.
